# TFBSpedia: a comprehensive human and mouse transcription factor binding sites database

**DOI:** 10.64898/2026.03.04.709638

**Authors:** Shiting Li, Elysia Chou, Kai Wang, Alan P. Boyle, Maureen A. Sartor

**Author notes:** Corresponding Authors: Maureen A. Sartor, University of Michigan, 1301 Catherine St Building 1, Ann Arbor, MI 48109.

## Abstract

Mapping the genomic locations and patterns of transcription factor binding sites (TFBS) is essential for understanding gene regulation and advancing treatments for diseases driven by DNA modifications, including epigenetic changes and sequence variants. Although several TFBS databases exist, no study has systematically benchmarked these databases across different sequencing technologies and computational algorithms. In this study, we addressed this gap by constructing a TFBS database that integrates all available ENCODE cell line ATAC-seq and Cistrome Data Browser ChIP-seq datasets, comprising 11.3 million human and 1.87 million mouse TFBS. We also integrated previously published TFBS resources (Factorbook, Unibind, RegulomeDB, and ENCODE_footprint) and found each contains a substantial fraction of unique TFBS predictions, highlighting significant discrepancies among existing resources. To assess the accuracy of the combined TFBS regions, we assembled ten independent genomic annotation datasets for evaluation and found that TFBS regions predicted by multiple databases are more likely to represent true and biologically meaningful binding sites. For each predicted TFBS region, we define two scores: the confidence score reflects prediction reliability, while the importance score represents biological functional relevance. Finally, we introduce TFBSpedia, a lightweight and efficient search engine that enables rapid retrieval of TFBS regions and comprehensive annotation information across the integrated databases.

## Introduction

Transcription factors (TFs) are proteins that bind specific DNA sequences and function as activators (or repressors) of gene expression ^1^. For *Homo sapiens*, approximately 1600 TFs have been identified, whereas *Mus musculus* possess ∼1200 known such proteins ^1,2^. Typically, TFs cooperate with other proteins such as RNA polymerase II to assemble transcriptional complexes near promoters or enhancers. These complexes then bind specific DNA regions, which are known as transcription factor binding sites (TFBS) to mediate transcriptional activation or repression ^3–5^.

A TFBS for an individual TF typically spans 8 to 20 base pairs. For certain genomic regions such as superenhancers, these TFBS often exist at high density, overlapping or clustering in close proximity, comprising a few hundred base pairs of TFBS regions ^6,7^. Given that a large percentage of TFBS are sequence specific and evolutionarily conserved ^8,9^, both epigenetic DNA modifications (methylation and hydroxymethylation) and genetic variants (SNP, INDEL, and SV) have the potential to alter the TF-TFBS binding affinity ^10,11^. These binding affinity changes can alter transcriptional programs, ultimately manifesting as phenotypic changes, including evolution, developmental disorders, and complex diseases ^12,13^. A paradigmatic example in oncogenesis involves the transcription factor *MYC*; here, the SNP rs6983267 alters *TCF4* binding affinity at a *MYC* enhancer, triggering increased TF occupancy that drives *MYC* overexpression, thereby promoting tumor growth and immune evasion ^14,15^. Beyond individual loci, retrotransposon-mediated mobilization allows TFBS to propagate across the genome, with recent studies indicating that 20% of TFBS are embedded within transposable elements (TEs) ^16,17^.

The ability to map and predict TFBS across the genome has dramatically improved with high-throughput sequencing technologies. ChIP-seq and its derivative technologies such as CUT&RUN directly capture DNA fragments bound by a specific TF protein ^18,19^. These DNA fragments are sequenced, followed by peak calling and statistical methods to pinpoint the TFBS locations. This remains the gold standard for TFBS identification. Constrained by various biological circumstances and challenges of antibody design, and coupled with the emergence of chromatin accessibility profiling sequencing, researchers have increasingly pursued computational methods (footprinting) using DNase-seq and ATAC-seq to predict TFBS genome-wide across multiple TFs simultaneously^20–22^. However, footprinting prediction faces a limitation: different TFs share overlapping binding sites, making it difficult to unambiguously predict TFBS for specific TFs ^23^. Additionally, to summarize the TF binding preferences across different conditions, position weight matrices (PWMs)—commonly referred to as motifs—were created ^24^. The motif stores the nucleotide frequency patterns at each position within TFBS. While these motifs are widely adopted ^25,26^, they are largely restricted to short monomeric sequences (6-20 bp) and often fail to capture the combinatorial complexity of TF-TF interactions and multi-protein binding patterns ^8,27^.

Benefiting from large-scale projects like ENCODE^28^ and Cistrome^29^, which systematically generate and/or assemble ChIP-seq, DNase-seq, and ATAC-seq data, a number of TFBS databases have been developed ^30,31^. However, no study has yet systematically benchmarked these motif-derived TFBS across databases by examining their appearance frequency, technical biases, and biological significance. In this study, we address this gap by developing the most comprehensive TFBS database and assessing the functional importance of each predicted TFBS region. To achieve this, TFBS regions detected by different algorithms were first compared, revealing substantial algorithmic biases. A new TFBS database, named UM TFBS database, was developed and integrated with four previously published resources. The resulting databases were evaluated at both regional and single-nucleotide resolution using ten different genomic annotations, showing that TFBS shared across multiple databases are more likely to represent true binding sites. Lastly, two scores (importance and confidence scores) were defined as references for TFBS region filtering. To maximize usability of the database, we developed TFBSpedia (https://tfbspedia.dcmb.med.umich.edu/), a web portal enabling efficient searching and downloading of the TFBS region information.

## Methods

### Collection of publicly-available position weight matrices

All available position weight matrices (PWMs) were downloaded from the JASPAR RESTful API 2020 ^32^, Hocomoco v11 full mono meme ^33^ and HOMER2 ^34^ for both Homo sapiens (hg38) and Mus musculus (mm10).

We obtained 971 human PWMs and 229 mouse PWMs from JASPAR, and 668 human PWMs and 356 mouse PWMs from Hocomoco. An additional 417 PWMs were retrieved from HOMER2 for both species. (**Supplementary Table 1**).

### Collection of ChIP-seq and ATAC-seq data for the UM TFBS database

To build the UM TFBS database, we first retrieved hg38 and mm10 TF ChIP-seq peaks from the Cistrome Data Browser ^35^ and scanned them using FIMO from MEME Suite 5.4.1 ^36^ for instances of the collected PWMs from JASPAR2020 and Hocomocov11 or HOMER2 with its own specific format of PWMs. For TFs with hg38 ChIP-seq data but without matched public PWMs, we utilized MEME-ChIP from MEME Suite and the HOMER2 de novo method to predict their motifs (507 human TF experiments, **Supplementary Table 1**). To ensure the accuracy and robustness of the de novo motif discovery from ChIP-seq data, we selected the top 6000 peaks ranked by signal level for each experiment as the input for MEME-ChIP and HOMER2. For the MEME-ChIP procedure, we kept motif results significantly centrally distributed among ChIP-seq peaks as defined by CentriMo.

Since only a subset of TFs have ChIP-seq data and are limited to specific cell lines, to extend the UM TFBS database, we collected ENCODE human and mouse cell line ATAC-seq peaks and bam files (**Supplementary Table 2**). We applied footprinting using TOBIAS v0.13.3 ^37^ and identified the additional TFBS based on ATAC-seq. All Hocomoco, JASPAR and new MEME-chip motifs were used, and only TOBIAS-predicted bound TFBS regions were included.

The TFBS from ATAC-seq and ChIP-seq data were merged. After additional quality control checks (See below), we named the final collection the UM TFBS database, containing both the hg38 and mm10 versions.

### TFBS quality control by integrated analysis of ATAC-seq and ChIP-seq

Different computational algorithms (FIMO, TOBIAS, and HOMER2) and sequencing technologies (ChIP-seq and ATAC-seq) may each have their own unique biases in predicting TFBS. Thus, to further validate our newly introduced motifs and characterize these biases, we defined a metric termed “coverage similarity” to quantify these technical differences for TF-cell line pairs. Assuming A and B are the predicted TFBS in two sets, the coverage overlap scores between them are calculated as (A⋂B)/|A| and (A⋂B)/|B|, and the max coverage similarity is defined as (A⋂B)/min(|A|, |B|).

Based on the number of predicted regions and the maximum coverage similarities calculated, we first excluded the TFs with fewer than 1000 TFBS predictions across all algorithms and sequencing technologies. Secondly, we removed motifs lacking inter-algorithm consistency, which we defined as having maximum coverage similarity below 0.1 across all pairwise comparisons. We also manually curated and removed putative TFs that lacked DNA binding affinity based on UniProt ^38^ (**Supplementary table 3**).

### Assembly of public TFBS databases

In addition to our UM TFBS database, we also assembled the TFBS from four established resources: Factorbook ^31^ (hg38), Unibind ^30^ (hg38,mm10), RegulomeDB all TRACE calling ^39^ (hg38) and the comprehensive ENCODE footprint project ^38^ (hg38). We downloaded these databases’ TFBS derived from complementary sequencing approaches: ChIP-seq (Factorbook, Unibind), Representative DNase Hypersensitive Site (rDHS, Factorbook only), DNase-seq and ATAC-seq (ENCODE, RegulomeDB); all datasets represent *in vivo* TFBS.

As Factorbook lacked the TF name for each TFBS, we cross-referenced the TFBS with ENCODE footprint motif scanning results ^39^. We tracked the genomic coordinates, cell types, and tissues and predicted TF gene symbols for all TFBS from all sources.

### Comparative analysis of the TFBS databases

We performed an overlap analysis to evaluate the concordance of TFBS predictions across five databases. Given the clustering of TFBS for different TFs within gene regulatory regions ^40^, we merged overlapping TFBS into TFBS clusters and then trimmed or expanded all clusters to 100bp (human) or 50bp (mouse) for each database separately. We then checked these overlapping TFBS regions using the *GenomicRanges findOverlaps* function and visualized them as upset plots using ComplexUpset v1.3.3 ^41^.

### Development of additional TFBS databases for benchmarking

We developed additional TFBS database versions to benchmark individual TFBS databases. For hg38, we built two complementary versions: *Union*, which merges all regions from the five TFBS databases using the *GenomicRanges reduce* function, providing the most comprehensive coverage; and *Intersection*, a more stringent version that includes only TFBS present in at least two independent databases. Because only two TFBS databases are available for the mouse mm10 genome, only the *Union* version was generated for evaluation. For additional annotations such as TF names, we collapsed them for each merged TFBS region.

### Annotation regions to benchmark TFBS databases

To evaluate the final assembled TFBS databases, we gathered six different types of genomic annotations for both human and mouse, and four additional annotations specific to human. These annotations include enhancers and promoters from Qin T, et al^42^, regulatory element-to-gene links (rE2Gs) across tissues ^43^, ENCODE candidate cis-regulatory elements (cCREs) ^44^, histone modification peaks associated with regulatory regions often bound by TFs (H3K4me1 which marks distal enhancers, H3K4me3 for active promoters, and H3K27ac for active enhancers/promoters) from Cistrome Data Browser ^29^, confident transposable elements (TE) from Dfam (bias<50 and e-value<10^-15^) ^46^, and ENCODE blacklist regions ^47^. For human-specific annotations, we incorporated all variants from the GWAS catalog V1.0.2 ^48^, tissue-specific expression quantitative trait loci (eQTL) V10 ^49^ and the most variable CpGs across tissues and cell types based on the DNA methylation level ^50^. Additionally, we collected all ChIP-seq and GHT-SELEX peaks from an independent TFBS database called Codebook ^51,52^.

### Regional and single base pair-level evaluation of TFBS databases

We calculated two parameters that represent the sensitivity and specificity of TFBS databases at both the regional level and single base pair (bp) resolution. Sensitivity is calculated as the percentage of evaluation regions that overlap with TFBS regions in databases, while specificity is defined as the percentage of TFBS regions that overlap with evaluation regions. We also calculated the Dice coefficient, which integrates both sensitivity and specificity, for single bp level evaluation. We quantified overlap at the regional level using the number of distinct regions, and at the single bp resolution using the number of base pairs. For single bp resolution evaluation, we also calculated rank-sum score for each TFBS database, assigning higher values to these with more overlap, except for the ENCODE blacklist, for which the scoring was reversed. Both evaluations and TFBS regions were collapsed with GenomicRanges as mentioned above to avoid duplicated regions being used. For evaluation regions normally longer than 2 bp (regions except GWAS, eQTL and variable CpGs), we required the overlap to be >5 bp for regional evaluation.

### Importance and confidence score calculations

We calculated two scores as evaluation metrics to assess the reliability and biological significance of predicted collapsed TFBS regions. The confidence score is calculated as the number of sources (five in total) and supporting sequencing techniques (ChIP-seq, ATAC-seq, and DNase-seq) for each predicted region, ranging from 0 to 8 for human and 0 to 4 for mouse. The importance score is calculated as the number of overlapping evaluation annotations, ranging from 0 to 7 for human and 0 to 4 for mouse.

### Development of TFBSpedia website

We developed the TFBSpedia website for ease of database access. The website database, which stores the collapsed TFBS regions and annotations, was implemented using PostgreSQL, while the website interface was developed using the Django framework. The website code is available at https://github.com/shengzhulst/TFBSpedia_website.

## Results

### Distinct sequencing assays and computational models introduce systematic biases in TFBS collections

To expand upon currently available transcription factor binding site (TFBS) databases, we first rigorously created a new University of Michigan (UM) TFBS database for both human and mouse (**See methods, Figure 1A)**. We then used this database to explore the biases introduced by various factors in TFBS prediction (**See methods, Supplementary Figure 1A**). By implementing two widely accepted TFBS prediction programs, FIMO and HOMER2, on all Cistrome Data Browser ChIP-seq peaks and a footprinting program, TOBIAS, on all ENCODE ATAC-seq data, we predicted TFBS for 2,432 transcription factor (TF)-cell line pairs (**Supplementary Table S3**). We performed pairwise comparisons for FIMO versus HOMER2 (same ChIP-seq peaks: 2412 experiments) and FIMO versus TOBIAS (same position weight matrix (PWM): 512 experiments) (**Supplementary Table S3**). Using the coverage similarity scores (**See methods, Supplementary Figure 1, Supplementary Table S3**), we observed unexpectedly high inconsistencies between the methods. Regarding the number of predicted TFBS, FIMO predicted on average 43% fewer TFBS regions than HOMER2 (**Supplementary Figure 1B)** while ATAC-seq-based TOBIAS predicted 50% fewer regions than ChIP-seq-based FIMO (**Supplementary Figure 1C**). We found that 32% of the TF-cell line pair comparisons had maximum coverage similarity (**See methods**) below 0.5 after strict quality control. Intuitively, this measure captures the degree of ‘nestedness’ between two TFBS collections, here indicating a low fraction of the smaller dataset captured by the larger (**Supplementary Figure 1D, Supplementary Table S3)**. Additionally, 27% (649/2412) of the TF-cell line pairs using the same ChIP-seq data but different prediction algorithms (FIMO versus HOMER) and 20% (103/512) using the same PWMs but on different sequencing technologies (ATAC-seq using TOBIAS versus ChIP-seq using FIMO) had below 0.1 max coverage similarity **(Supplementary Table S3**). These low-level overlaps indicate that different sequencing technologies, algorithms, and motifs each have their own biases in predicting TFBSs even when using the same biological sources. To overcome this, we only included consistent predictions (i.e. TF-cell line pairs predictions have max coverage similarity larger than 0.1) in the UM TFBS database to increase its specificity (**See Methods, Supplementary Figure 1A**).

**Figure 1.**
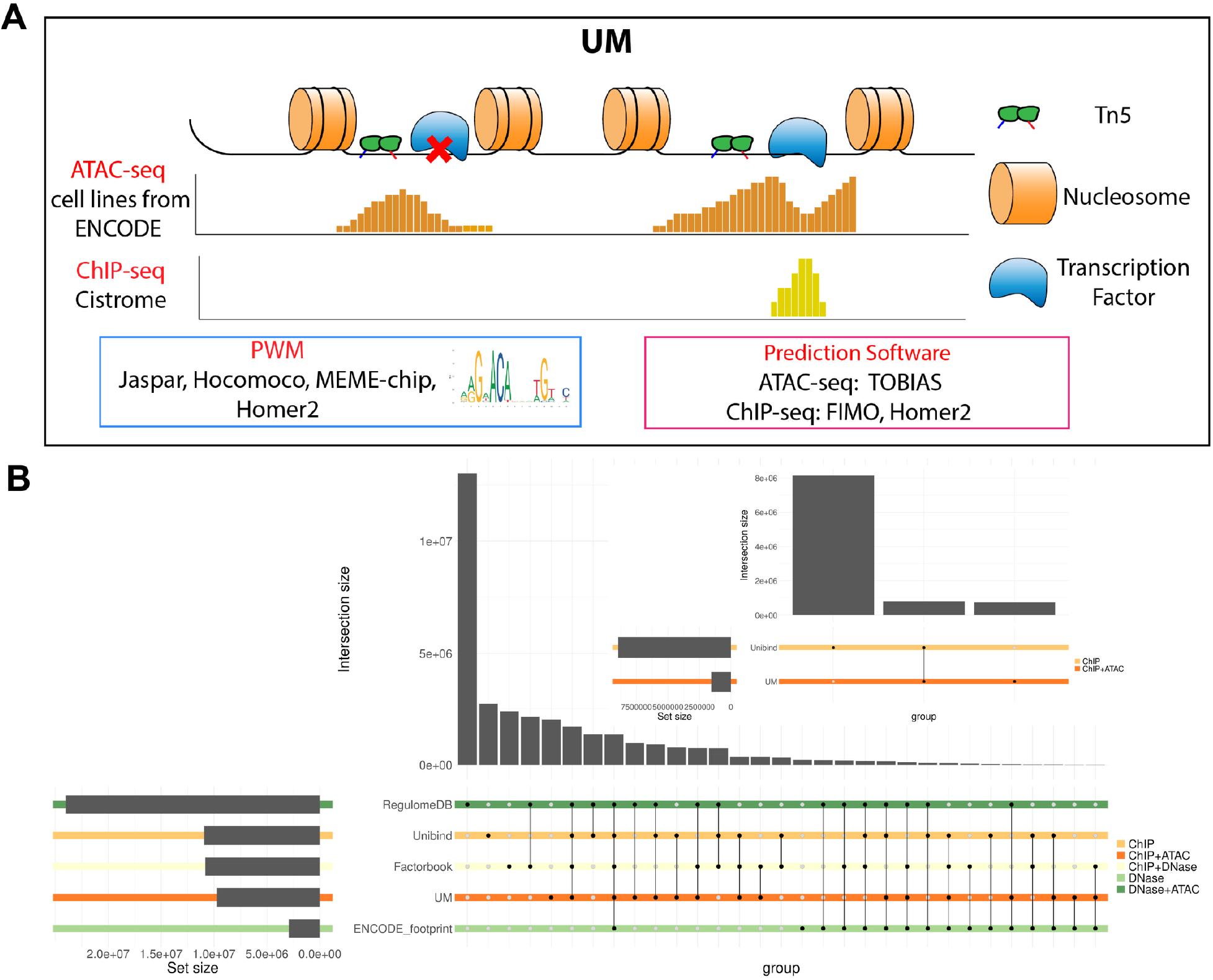
Overview of UM TFBS database creation and cross-database benchmarking. (A) Workflow schematic describes the data sources and computational software used to generate UM TFBS database. (B) Upset plots (main panel: hg38; top-right panel: mm10) comparing UM TFBS with four published TFBS databases (RegulomeDB, UniBind, Factorbook and ENCODE_footprint), colored by combinations of sequencing technologies.

To evaluate the newly created UM database, we benchmarked it against four published TFBS databases (Unibind, Factorbook, Encode Footprint and RegulomeDB) (**Table 1**). Only the UM and UniBind TFBS databases included mouse. We collapsed or expended the TFBS regions into 100 bp (human) and 50 bp (mouse) windows for an unbiased comparison (**See methods**). By examining the overlaps among these databases, we found that each contains a substantial number of unique regions for both hg38 and mm10 (**Figure 1B**). For hg38, the total number of TFBS regions and the number of unique regions followed the same descending order: RegulomeDB, UniBind, Factorbook, UM, and ENCODE_footprint, with RegulomeDB and Factorbook showing extensive regions overlapped with each other, likely derived from them both using DNase-seq footprinting (**Figure 1B**). These results illustrate that each of the current individual TFBS databases is likely insufficient both in terms of coverage and accuracy; integrating them may provide higher quality TFBS region information.

**Table 1.** Description of the public available TFBS databases.

| Database name | Sequencing Tech | Sequencing and Peak sources | TFBS prediction software | Motif sources (database or software) | Year published |
| --- | --- | --- | --- | --- | --- |
| ENCODE_footprint | DNase-seq | ENCODE | empirical Bayes framework | Taipale, JASPAR, HOCOMOCO | 2020 |
| Unibind | ChIP-seq | Remap, GTRD | ChIP-eat | JASPAR, DEMO | 2021 |
| Factorbook | ChIP-seq, DNase-seq | ENCODE | FIMO | Factorbook | 2022 |
| RegulomeDB | DNase-seq, ATAC-seq | ENCODE | TRACE | ENCODE | 2023 |
| UM | ATAC-seq, ChIP-seq | ENCODE, Cistrome Data Browser | FIMO, TOBIAS | JASPAR, HOCOMOCO, HOMER2, MEME-CHIP |  |
| Cookbook | ChIP-seq, GHT-SELEX | Cookbook |  |  |  |

### Benchmarking TFBS databases by multiomics evaluations

To further compare and quantify the differences among these available TFBS databases and create a more comprehensive and accurate TFBS database, we created two additional TFBS collections based on the union and intersection of at least two of the existing databases, named Union and Intersection (**Figure 2A, see methods**). To evaluate the TFBS databases, we collected ten different genomic annotations for evaluation (**Figure 2B, see methods**), covering widely conceptualized gene regulatory elements including (i) promoters, (ii) enhancers, (iii) histone modifications associated with active enhancers or promoters (H3K4me1 H3K4me3, and H3K27ac), (iv) transposable elements (TE), ENCODE annotations including (v) cCREs, (vi) significant variants from GWAS, (vii) eQTLs from GTEx, (viii) variable CpGs, defined as CpGs with high methylation variability across tissues and cell types and potentially linked to cell differentiation, (ix) the ENCODE blacklist, and (x) a recently introduced rare TFs’ TFBS database (Codebook). We hypothesized that a higher-quality TFBS database would show greater overlap with these independent DNA annotations while covering less of the ENCODE blacklist.

**Figure 2.**
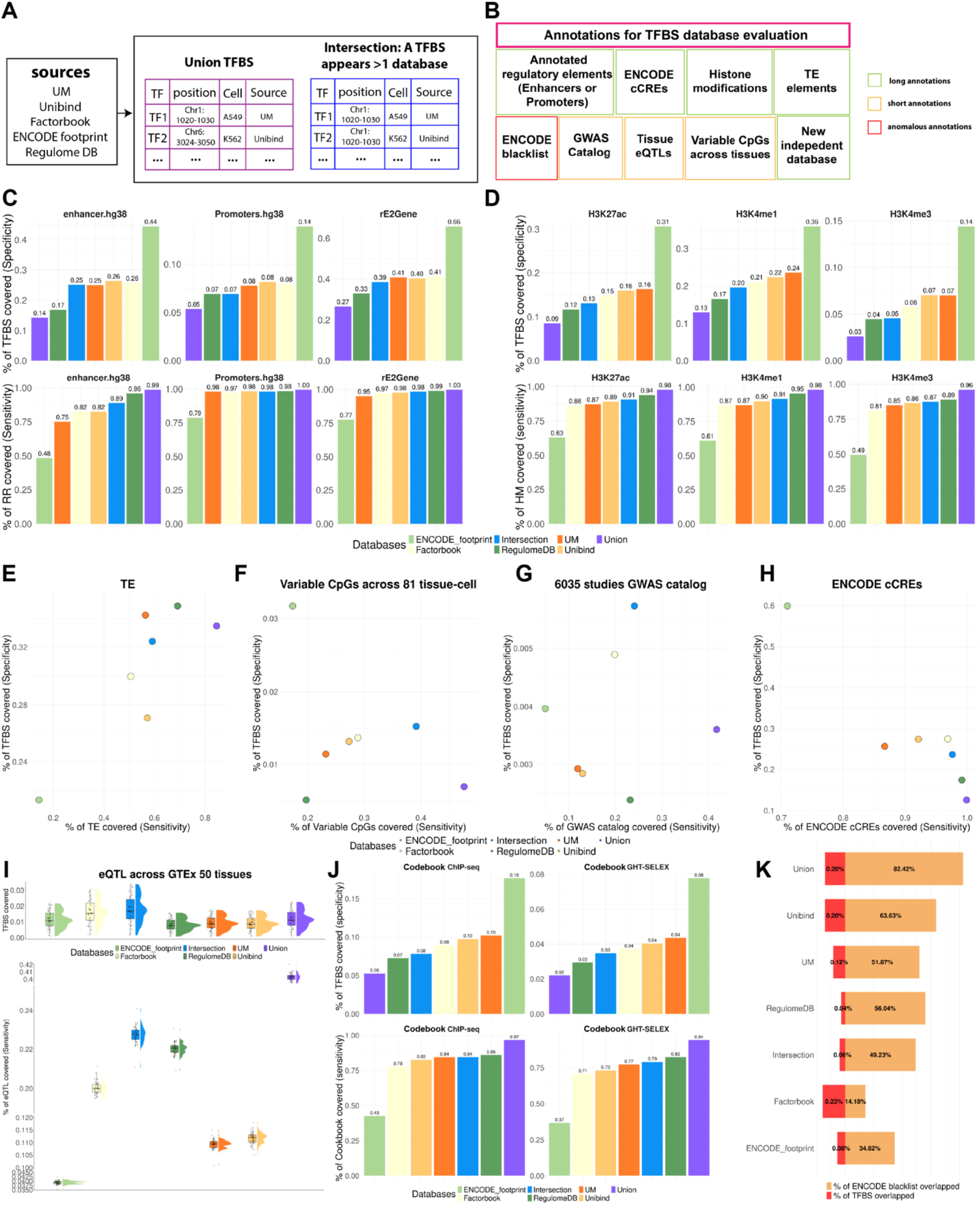
Regional-level evaluation of TFBS databases and two new integrated TFBS collections for hg38. (A) Description of the two new TFBS collections derived from existing TFBS databases. (B) Overview of the genomic annotation sets used to benchmark TFBS databases. (C-J) Bar, dot and raincloud plots summarizing the sensitivity and specificity of each TFBS database and set across annotation sets, including (C) promoters, enhancers, (D) transcription-related histone modification peaks, (E) TE, (F) variable CpGs, (G) GWAS catalog, (H) ENCODE cCREs, (I) eQTLs from GTEx, (J) ChIP_seq and GHT-SELEX peaks from Codebook. (K) Sensitivity and specificity evaluated against the ENCODE blacklist, lower overlap indicates higher database quality.

We first examined the genomic coverage of the TFBS databases and the collected genomic annotations for evaluation. For both human and mouse, the genomic annotations together covered ∼50% of the relevant genome, with TEs (25-30% of all genomes) tending to be less overlapped with others (**Supplementary Figure 2A, D)**. We collapsed single TFBS into larger TFBS regions to simplify the analysis due to the extremely high level of TFBS overlapping (**See methods**). For example, within a 124-bp enhancer region: chr1: 940453-940577, we observed potential binding from 162 different TFs across 68 cell lines/tissues based on all databases. All predicted TFBS regions from all databases (Union) cover nearly 30% of the human genome and 7% of the mouse genome, with the Intersection set (i.e. occurring in at least two databases) covering 15% of the human genome (**Supplementary Figure 2B, E**). 97% of the collapsed TFBS regions in the Union set were shorter than 100 bp, with Union and RegulomeDB having nearly 30 million TFBS regions and others having between 11 and 13 million. The exception was ENCODE footprint, which only contains ∼3.5 million human TFBS regions (**Supplementary Figure 2C, F**).

We next examined the overlap between the TFBS databases and the evaluation genomic annotations (**Figure 2C-K**). We defined the *specificity* of each TFBS database as the percentage of the TFBS regions overlapped and the *sensitivity* as the percentage of the evaluated regions overlapped (**see methods**). The optimal TFBS database should have higher values of both, except for the ENCODE blacklist. For humans, as expected, the Union set consistently had the highest sensitivity and lowest specificity, and that ENCODE_footprint, which only included TFBS appearing in multiple cell lines, consistently had the highest specificity and lowest sensitivity (**Figure 2C-J**). Specifically, for specific gene regulatory elements (rE2G enhancers, promoters, and histone modifications), each database overlaps ∼90% or more of these annotations, and the Intersection performed similarly to RegulomeDB (**Figure 2C-D**). RegulomeDB outperformed in terms of capturing TEs and another ENCODE annotation, cCRE, and the Intersection set demonstrated the best performance for all short region annotations (variable CpGs, GWAS and eQTLs) (**Figure 2E-I**). Although the independent Codebook project targeted 332 putative TFs not characterized in any other TFBS database, the Union set nonetheless covered 97% and 94% of their ChIP-seq and GHT-SELEX peaks, respectively, indicating its broad and comprehensive coverage (**Figure 2J**). All individual TFBS databases and our generated Union and Intersection sets have less than 0.25% of regions overlapped with the ENCODE blacklist, suggesting their robustness to technical issues (**Figure 2K**). For mouse, which has only Unibind and the UM database, Unibind was six times larger than the UM database (**Supplementary Figure 2E)**, and the UM database did not contribute much to increasing the TFBS region sizes (**Supplementary Figure 2F**) or coverage of the genomic annotations for evaluation (**Supplementary Figure 3**).

As the TFBS regions have different genomic lengths compared to the genomic annotations for evaluation, calculating the number of overlapped regions without considering the number of base pairs overlapped between each pair of regions is potentially biased. Therefore, we further benchmarked these TFBS databases at the single bp level for more accurate evaluation (**see methods**). To balance specificity and sensitivity, we calculated Dice coefficients for the TFBS databases and sets and an overall rank sum for each TFBS database. Based on Dice coefficients, the Intersection set ranked best, followed by the UM database second (**Figure 3A-C, Supplementary Table S4**). Unlike the region-level evaluation, the Intersection set clearly outperformed RegulomeDB and all other individual TFBS databases in sensitivity (**Figure 3D-F**), and closely followed Unbind and the UM database in terms of specificity (**Figure 3G-I**). All single bp level evaluations demonstrated that TFBS regions present in multiple databases tend to be in biologically meaningful regions.

**Figure 3.**
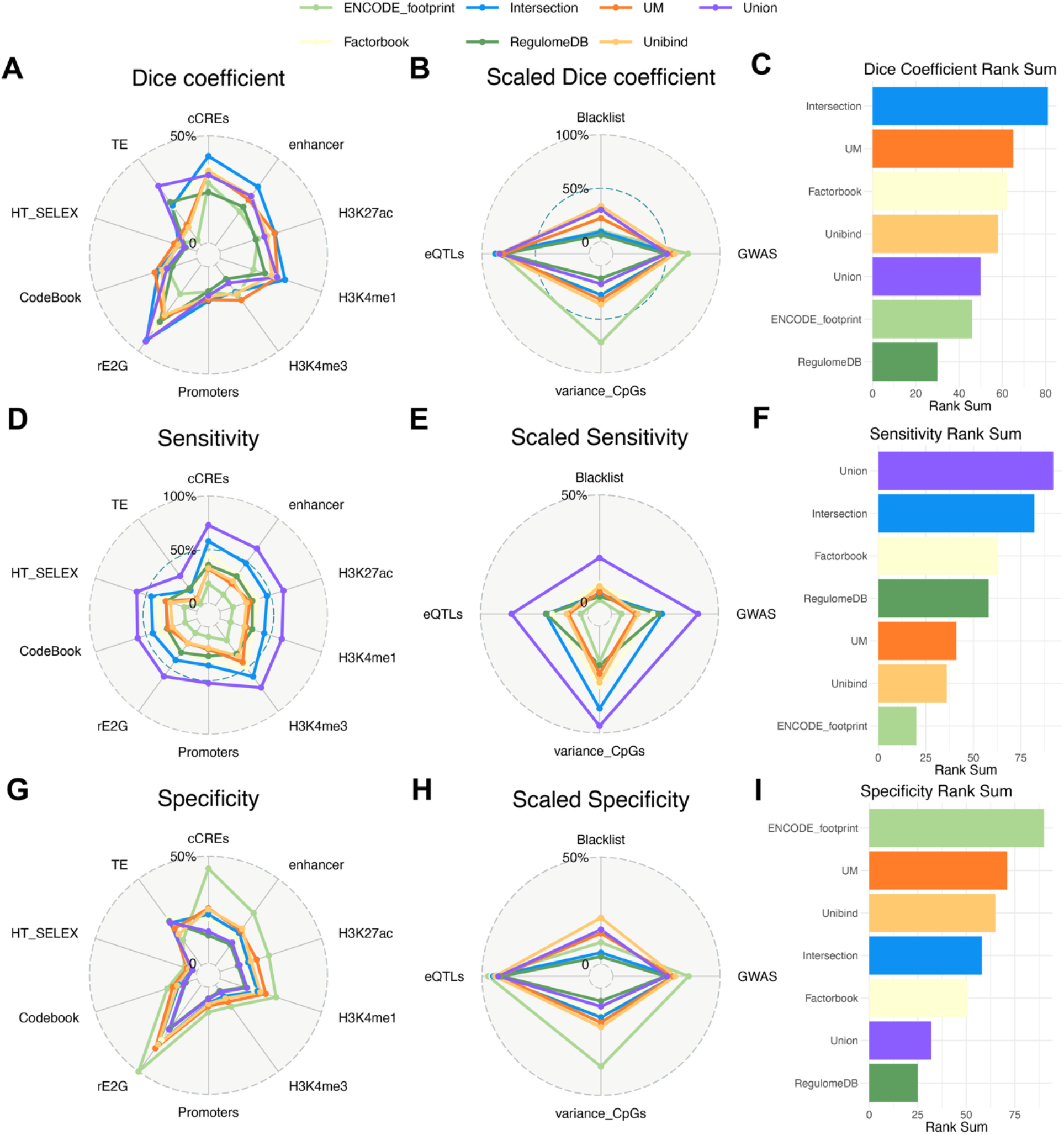
Single-base-pair (bp) level evaluation of TFBS databases and two new integrated TFBS collections. (A-I) Radar and bar plots summarize the performance using dice coefficient, sensitivity and specificity, along with metrics summarized by rank-sum scoring. (A-C) Dice coefficient, (D-F) Sensitivity (G-I) Specificity.

### TFBS region scores and web server development

As shown above, TFBS regions that appeared in at least two databases are more likely to be in other regulatory-associated annotations; thus, we have more confidence in them. Similarly, we reasoned that these regions are more biologically important. To systematically present this information for all TFBS regions, we defined two complementary scores, a *confidence* score and an *importance* score for each TFBS region. The *confidence* score represents the likelihood of a TFBS region being captured by five different TFBS databases and technologies (ATAC-seq, DNAse-seq, or ChIP-seq). The *importance* score represents the level of biological significance of each TFBS region, defined as the number of overlapping genomic annotations minus one if it overlaps with the Blacklist (**see methods**). TFBS regions appearing in multiple databases and sequencing technologies are potentially more frequently bound by TFs across conditions. As expected, the confidence scores and importance scores were correlated (Spearman rank’s correlation coefficient: 0.16 for mouse and 0.26 for human, p-value<2.2e-16 for both; **Figure 4A**). Furthermore, the importance scores significantly correlated with the number of cells/tissues in which a TFBS region was detected (**Figure 4B**). These results support our scoring framework and demonstrate that TFBS regions predicted recurrently across different conditions are more likely to play critical biological functions.

**Figure 4.**
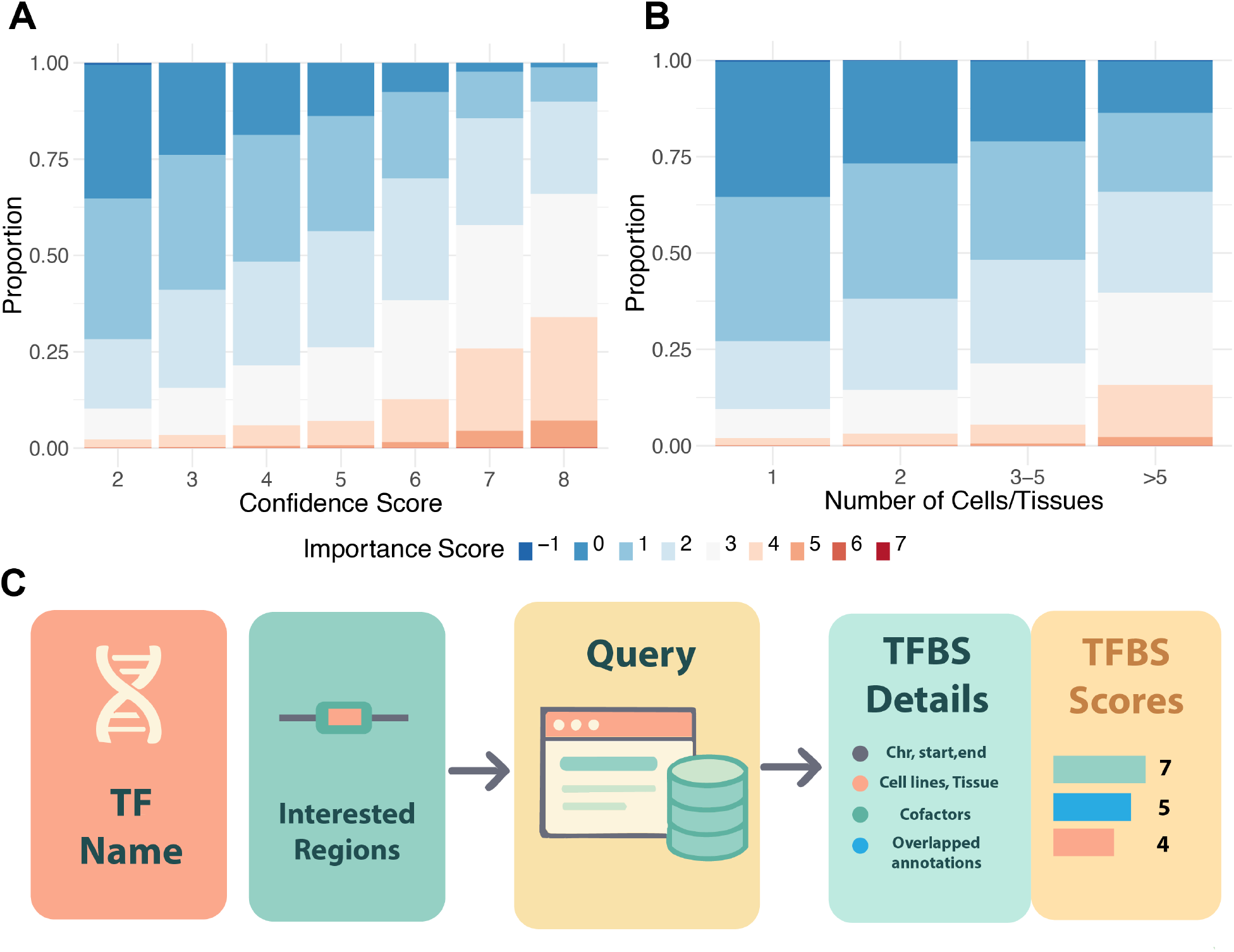
Confidence and importance scores and overview of TFBSpedia web portal. (A) Stacked bar plot showing the correlation between confidence score and importance score across TFBS regions. (B) Stacked bar plot showing the correlation between importance score and the number of cell lines/tissues in which a TFBS regions is detected. (C) Schematic overview of the TFBSpedia website and its query/annotation outputs.

Lastly, we developed a user-friendly website https://tfbspedia.dcmb.med.umich.edu/ for easy retrieval of TFBS region information. Users may query by TF gene symbol or genomic regions of interest; results contain the filtering and selection details and scores for each potential TFBS region (**Figure 4C**).

## Discussion

Current transcription factor binding site (TFBS) databases have limited coverage across diverse biological conditions, and are biased by the specific sequencing technologies utilized. Because users are often interested in specific SNPs, we performed evaluations and provide annotations at the single bp level. In terms of sequencing technologies, chromatin accessibility-based methods like DNase-seq and ATAC-seq exclusively target open chromatin binding sites, consequently missing some transcription factor binding sites in unopened chromosomal regions, especially for pioneer TFs such as those of the FOXA family ^53–55^. Additionally, existing databases are affected by computational framework biases, as differences in peak-calling algorithms and processing pipelines can substantially influence binding site identification, as shown in our results. To address these limitations, we developed TFBSpedia, by integrating all available TF binding sites from the three primary experimental assays (ChIP-seq, ATAC-seq, and DNase-seq), analyzed using different computational protocols, effectively expanding the coverage of known TFBSs in human and mouse. Rather than focusing on individual TF motifs, TFBSpedia implements a genomic region-centric strategy that systematically targets TFBS across the entire genome while still providing TF motif information and annotating them with extensive independent genomic annotations. By collecting the most comprehensive assembly of TFBS regions to date, together with a novel two-score framework that quantifies the confidence and importance of each TFBS region, TFBSpedia will serve as a new gold standard for reference genomic annotations.

As demonstrated in previous studies, TFBS regions are densely concentrated within specific gene regulatory elements, such as super-enhancers, where their spatial orientation and organization play a critical role in modulating regulatory activity and transcriptional output ^56^. Consequently, our analysis found that certain genomic regions are dense with overlapping TFBSs, with some of these regions extending up to several hundred bp. These complex, multi-TFBS regions have been shown to contribute strongly to gene regulation ^7,54–56^. To help us learn these TFBS regions, we provide evaluation scores and comprehensive annotations for these regions that are absent in existing TFBS databases. Additionally, our approach enables researchers to prioritize regulatory regions for experimental validation based on our score systems and to disentangle the cooperative mechanisms underlying transcriptional control.

Despite its many strengths, TFBSpedia has a few limitations. The most significant limitation is its reliance on predefined motifs from public motif sources, which are lacking for some not well-studied TFs. such as some zinc finger TFs ^57^, or binding sites with high sequence variability ^58^. Alternative deep learning methods, like ChromBPNet ^59^, use neural networks for footprints, providing more flexibility in defining TF syntax and accommodating sequence variants. However, our benchmarking analysis shows that TFBSpedia covers 95% of the new independent dataset Codebook which targets rare understudied -TFs, providing evidence that TFBSpedia has excellent genomic coverage across a wide range of conditions. Future efforts should focus on comprehensively characterizing how defined TF binding activity at specific sites varies across cellular contexts, developmental stages, and diseases.

