## Supplementary figures and table legends for "TFBSpedia: a comprehensive human and mouse transcription factor binding sites database"

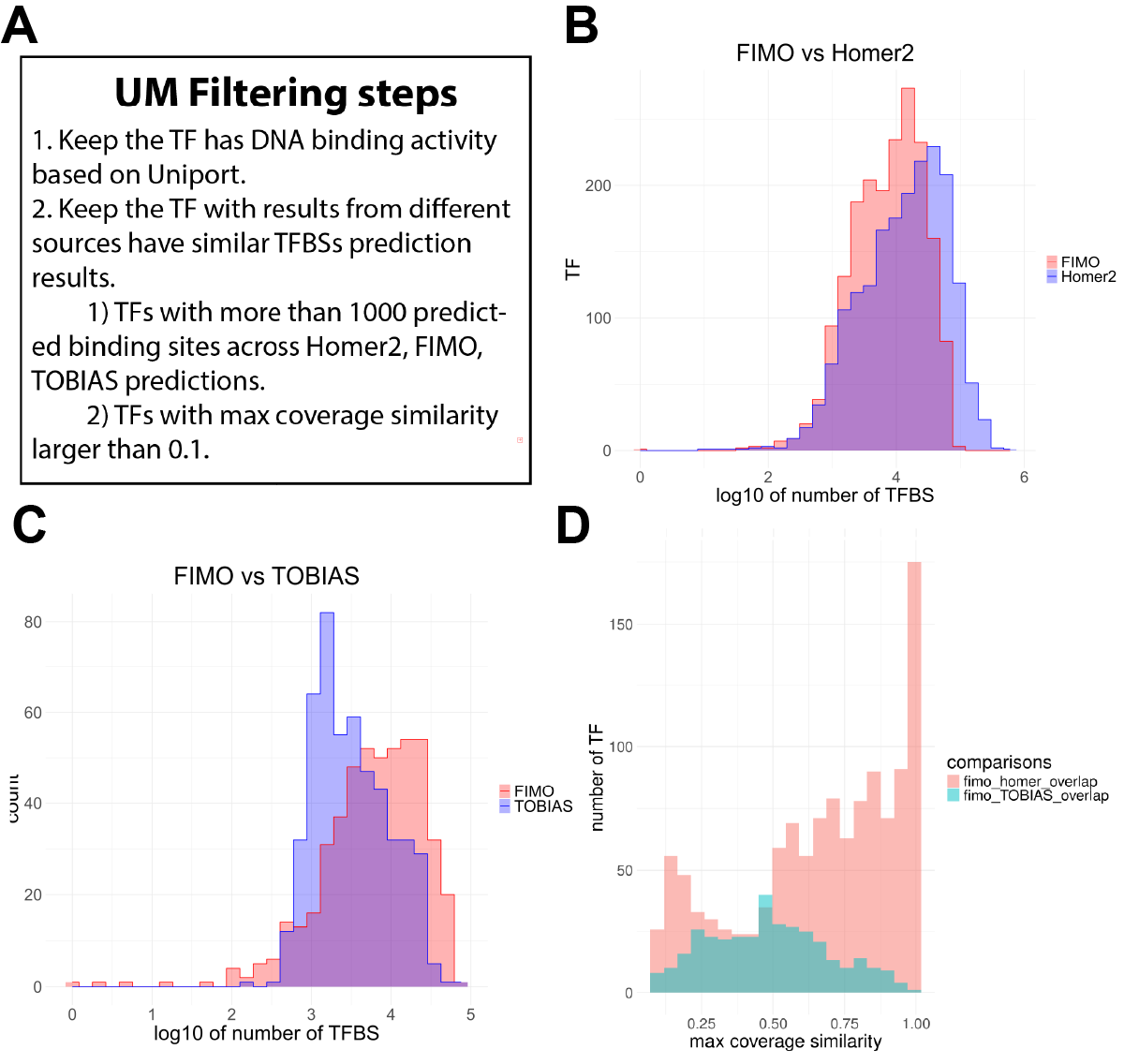

**Supplementary Figure S1. Workflow for filtering the UM TFBS database and comparison across TFBS prediction algorithms** (A) Filtering steps used to obtain high quality TFBS. (B-C) Comparison of the number of TFBS predictions generated from different algorithms using matched cell line/tissue-TF pairs. (D) Comparisons of max coverage similarly between fimo and homer (different prediction algorithms) or between fimo and TOBIAS (sequencing technology differences).

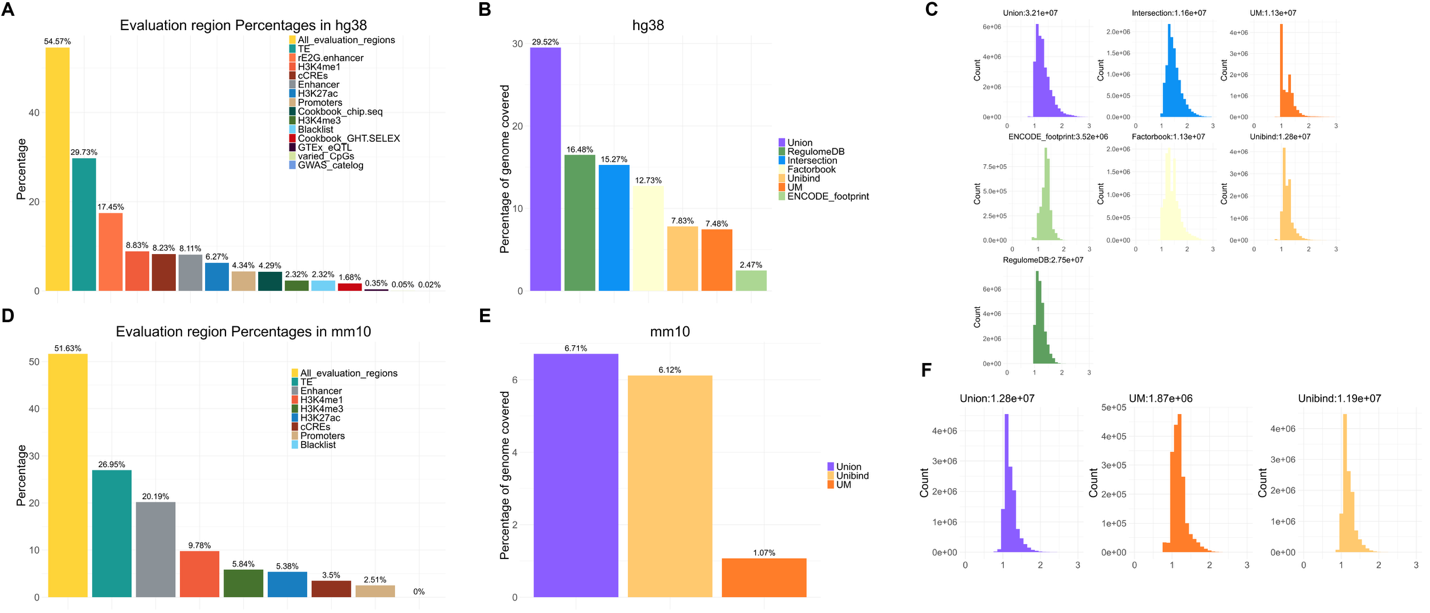

**Supplementary Figure S2. Genomic coverage and length distribution of TFBS databases and collections and annotation sets** (A) Evaluation genomic annotation coverage in hg38. (B) TFBS databases and collections coverage in hg38. (C) The length distribution of collapsed TFBS regions across databases and collections in hg38 (D-E) Genomic annotations, TFBS databases and collections coverage for mm10. (F) Length distribution of collapsed TFBS regions across databases and collections in mm10.

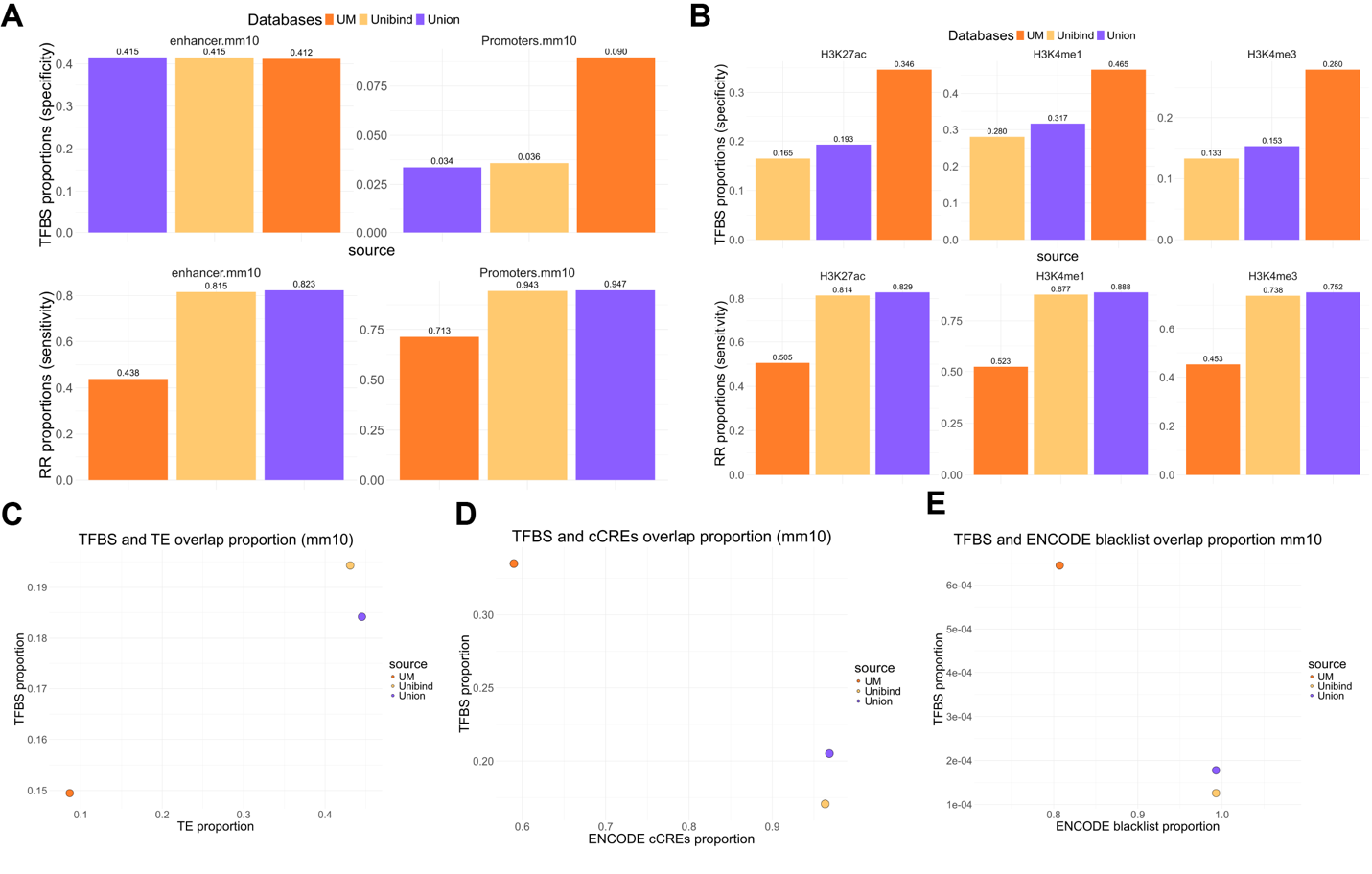

**Supplementary Figure S3. Regional-level evaluation of TFBS databases and one new integrated TFBS collections for mm10.** Bar and dot plots summarizing the sensitivity and specificity of each TFBS database and set across annotation sets, including (A) promoters, enhancers, (B) transcription-related histone modification peaks, (C) TE, (D) ENCODE cCREs, (E) Sensitivity and specificity evaluated against the ENCODE blacklist, lower overlap indicates higher database quality.

**Supplementary Table S1.** Transcription factor motif sources across databases. Motifs collected from JASPAR (human and mouse), HOCOMOCO (human and mouse), HOMER 2, and de novo approaches.

**Supplementary Table S2.** ENCODE ATAC-seq file accessions with species and cell line information used for UM TFBS database.

**Supplementary Table S3.** Comparison of FIMO, HOMER and TOBIAS predicted TFBS regions and calculated overlap coverage per TF-cell line combination.

**Supplementary Table S4.** Single bp level evaluation of TFBS databases and collections across sources using sensitivity, specificity, and dice coefficient.
